# Covariance-based decoding reveals content-specific feature integration and top-down processing during visual imagery

**DOI:** 10.1101/2022.09.26.509536

**Authors:** Francesco Mantegna, Emanuele Olivetti, Philipp Schwedhelm, Daniel Baldauf

## Abstract

When we internally generate mental images, we need to combine multiple features into a whole. Direct evidence for such feature integration during visual imagery is still lacking. Moreover, cognitive control mechanisms, including memory and attention, exert top-down influences on the perceptual system during mental images generation. However, it is unclear whether such top-down processing is content-specific or not. Feature integration and top-down processing involve short-range connectivity within visual areas, and long-range connectivity between control and visual areas, respectively. Here, we used a minimally constrained experimental paradigm wherein imagery categories were prompted using visual word cues only, and we decoded face versus place imagery based on their underlying connectivity patterns. Our results show that face and place imagery can be decoded from both short-range and long-range connections. These findings suggest that feature integration does not require an external stimulus but occurs also for purely internally generated images. Furthermore, control and visual areas exchange information specifically tailored to imagery content.

**Teaser:** Decoding visual imagery from brain connectivity reveals a content-specific interconnected neural code for internal image generation.

## Introduction

Our brain has a remarkable capacity to internally generate vivid mental representations in the complete absence of external sensory stimulation. Imagery is very useful whenever we need to process information that is not accessible in the present. For example, imagery allows us to re-instantiate information encountered in the past or to anticipate information that we will encounter in the future, without constantly requiring an external reference. In particular, visual imagery involves the internal generation of mental images ^1^. We can generate rich, vivid, and detailed images in our mind’s eye, which can contain precise color and shape information. For example, we can internally visualize a well-known place or a familiar person’s face. In both cases, the imagined percept may involve visual details with particular shapes, colors, hues, textures, and shading. This implies that multiple visual features have to be integrated to yield a complex and coherent visual representation ^2^.

Feature integration requires communication between brain areas that locally represent specific visual features of the imaginandum. Various areas need to communicate in order to combine dispersed feature representations into a coherent visual percept. Information integration is at the core of a cortical processing model proposed by Tononi and colleagues ^3^. This model is supported by computer simulations suggesting that reciprocal information exchange across areas in the visual cortex is the basic computational mechanism for information integration ^4^ This computation is biologically plausible insofar as areas in the visual cortex are strongly interconnected ^5^. There is also empirical evidence for the fact that patterns of neuronal synchronization reflect information integration during visual perception ^6^. However, it is unknown whether the same feature integration mechanisms are also employed when generating mental images internally, in the absence of any sensory stimulation. Neuroimaging studies have shown that areas that locally represent specific features during visual perception also represent the same features during visual imagery. For example, the primary visual cortex represents spatiotopic information ^7^, area MT represents motion ^8^, while specialized areas in the inferior-temporal cortex represent faces and places, respectively ^9^. Therefore, the cross-talk between visual areas may be the very basis of complex mental image formation.

Short-range connections between visual areas alone are presumably not sufficient to achieve feature integration when there is no external stimulus. Instead, visual areas may be supported by cognitive control mechanisms, such as memory and attention, exerting top-down influences during imagery, as suggested in the model proposed by Sakai and Miyashita ^10^. In particular, their model suggests that different objects or parts of a scene must be retrieved from a memory storage and are visualized using focal attention during imagery. This model is supported also by neuroimaging evidence suggesting that not only visual areas but also frontal and parietal areas associated with top-down processing are activated during imagery ^11^. There is also evidence that occipital and temporal areas receive top-down inputs from frontal and parietal areas through long-range connections during imagery ^12^. However, it is unknown whether top-down signals are specific to different imagery targets (e.g., faces versus places) or not. Functional connectivity patterns can be specific in strength, spatial destinations, or both. Watrous and colleagues ^13^ have shown that connectivity strength between temporal, parietal and frontal areas during visual perception is associated with better subsequent spatio-temporal memory retrieval. Moreover, Baldauf and Desimone ^14^ observed connectivity patterns having specific spatial destinations - from the inferior frontal junction to either the fusiform face area or the parahippocampal place area - depending on whether participants were instructed to pay attention to faces or places during visual perception. Collectively, these studies suggest that top-down signals exerted through long-range connections may vary also depending on the content of visual imagery.

Neural decoding is an excellent tool to address questions about content-specific representations ^15^. It uses machine learning algorithms to read out different stimulus categories from brain signals. Content specific information has been successfully decoded during visual perception, for example, by deciphering single visual features (e.g., orientation, shape, color) but also more complex object information from recorded brain signals ^16, 17, 18^. Similar approaches have been used to try to decode visual imagery, too. For instance, previous studies tried to decode different types of content specific information (e.g., perceptual, conceptual) during visual imagery from the sparse activation of various brain areas over time ^19, 20,21^.

In this magnetoencephalography (MEG) study, we test the hypothesis that different imagery categories are associated with distinct functional connectivity patterns reflecting content-specific feature integration and top-down processing. Participants were asked to imagine two different types of targets: faces and places. In contrast to previous studies, we instructed the two imagery categories by using word cues only, rather than showing any concrete pictorial aids. The rationale was that - in the absence of any pictorial references - participants have to internally generate mental images purely based on memory and attentional control. Consequently, any differences between imagery categories would be fully attributable to an internally driven effort to (re-)instantiate mental images rather than being confounded with low-level visual information artificially introduced by a pictorial aid. We hypothesized that the imagery of faces and places involves distinct feature integration and top-down processes, associated with distinct connectivity patterns of different strength, spatial destination, or both. To test this hypothesis, we used neural decoding to read out imagined categories from the connectivity patterns measured across MEG sensors as well as the reconstructed cortical sources. To achieve that goal, we used a connectivity decoding method based on spatial covariance that was originally applied to motor imagery for brain computer interface (BCI) applications ^22^. This decoding method was designed to capitalize on relative changes in brain activity measured from M/EEG sensor pairs. Here, we tested whether it will allow us to capture connectivity patterns distinguishing face versus place imagery, i.e. a type of internal signal that is notoriously hard to decode with classic time-domain decoders due to the temporal misalignment across trials. Moreover, in order to disentangle the contributions of feature integration and top-down processing, we test to what extent the decoding is driven by short-range and long-range connections.

## Results

### Decoding performance evaluation on simulated data

To test our predictions we scanned participants using magnetoencephalography (MEG) while they performed a visual imagery task. The task (Fig. 1A) consisted in the internal generation of a mental image of either a face or a place, randomly intermixed and cued by a word cue on a trial-by-trial basis. Participants were instructed to imagine a familiar instance of the cued category for several seconds while keeping their eyes fixated at the center of the screen. At the end of each trial, participants had to rate the vividness of their imagination and to confirm the imagined category of the present trial.

**Figure 1.**
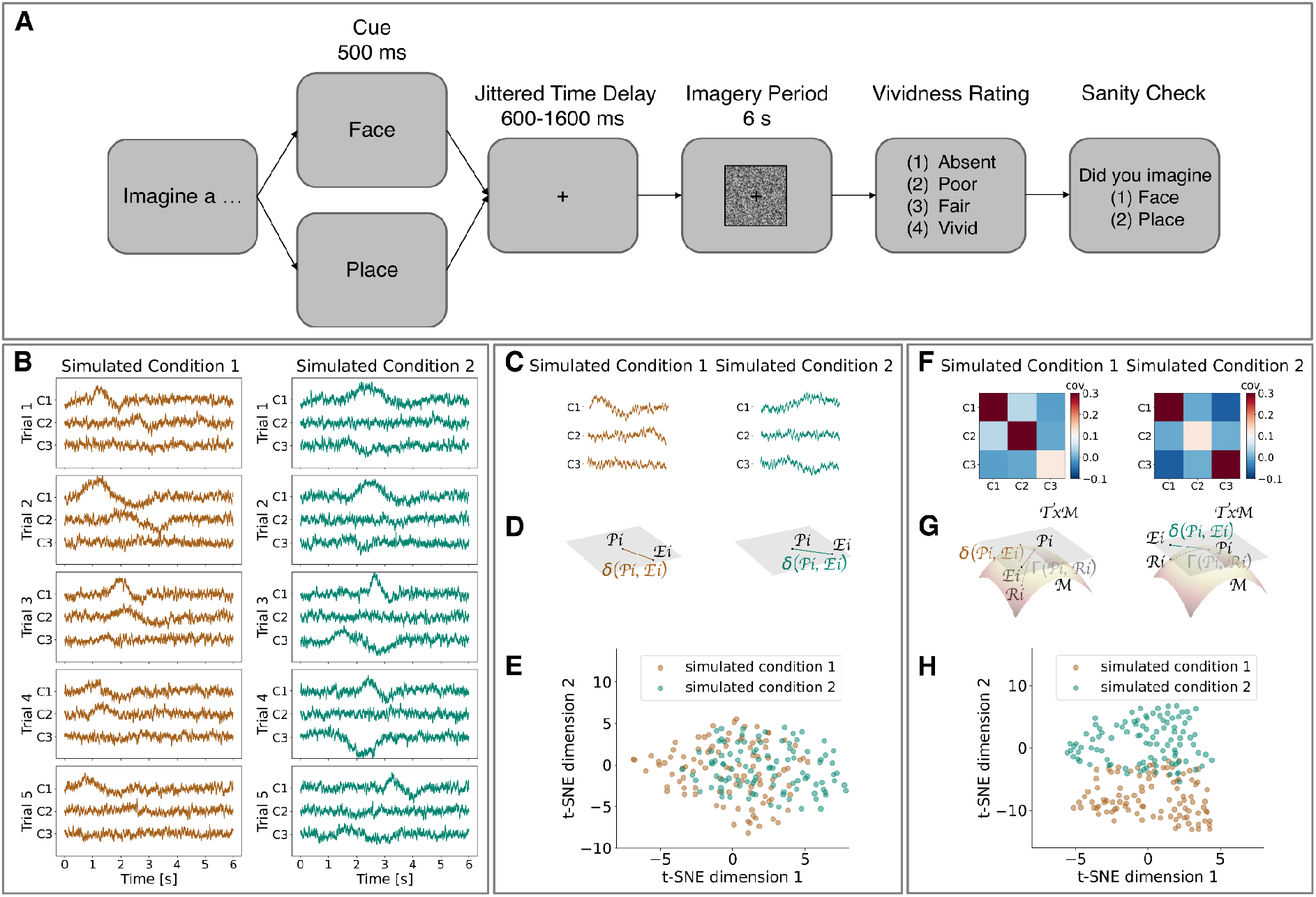
Experimental procedure, decoding methods and simulated data. The experimental procedure **(A)** was structured as follows: each trial began with a visual word cue instructing one category as the imagery target; then, a jittered time delay ensued after which the subject had to imagine a familiar instance of the cued category while a dynamic phase-scrambled mask was presented on the screen for 6s. At the end of each trial, participants were asked to rate the vividness of their imagination and confirm the category they had imagined during the trial. In a computer simulation **(B-H)** we tested whether relevant information is captured by covariance-based decoding or classic time-domain decoding. For this purpose, we simulated time series belonging to two different conditions **(B)** (here, we show only 5 representative trials in 3 simulated channels). Each trial contained a signal simultaneously embedded in noise of different channels. However, the onsets (and offsets) of the signal were misaligned across trials and across channels. To prepare input data for time-domain decoding **(C-E)**, we concatenated the time series of various channels into one single vector for each simulated condition **(C)**. The distance between an exemplar vector (*Ei*; i.e., single trial) and a prototype vector (*Pi*; i.e, average across trials) for each simulated condition can be estimated using an Euclidean metric (*δ*) **(D)**. Dimensionality reduction **(E)** of all exemplar vectors - corresponding to all simulated trials - shows that simulated conditions are not linearly separable when using time-domain decoding. To prepare input data for covariance-based decoding **(F-H)** we estimated spatial covariance matrices for each condition **(F)**, measuring the interdependence between channel pairs. The distance between an exemplar matrix (*Ri*) and a prototype matrix (*Pi*) on a Riemannian manifold (*M*) can be estimated using a Riemannian metric (*┌*). Then, the covariance matrix can be projected to an Euclidean tangent space (*TxM*) obtaining a tangent vector **(G)**. After that, the distance between an exemplar tangent vector (*Ei*; single trial) and a prototype tangent vector (*Pi*; average across trials) for each simulated condition can be estimated using an Euclidean metric (*δ*). Dimensionality reduction **(H)** of all tangent vectors - corresponding to all simulated trials - shows that simulated conditions are linearly separable when using covariance-based decoding.

First, we ran a simulation to investigate whether relevant information is captured by covariance-based decoding or classic time-domain decoding (see Fig.1B-H). There are major challenges associated with visual imagery signals - or, more in general, with any internally generated brain signal: On the one hand, information is temporally misaligned because there is a high variability in the onsets and offsets of imagination events across trials. On the other hand, information is presumably encoded in the reciprocal interconnections between channel pairs that give rise to specific spatial configurations. To account for these two aspects we simulated data as follows. We generated time series for one hundred trials in three different simulated channels. Each time series consisted of a combination of signal and noise. To account for temporal misalignment, we added random delays to signal onsets and offsets. To account for specific spatial configurations, we simulated data such that trials belonging to the first simulated condition had higher amplitude modulation in the first and the second channel while trials belonging to the second simulated condition had higher amplitude modulations in the first channel and the third channel (see Fig.1B). Then, we prepared input data for linear classification by using two different vectorization procedures. To prepare data for classic time-domain decoding, we concatenated time series corresponding to different channels into one single vector (Fig.1C). To prepare data for covariance-based decoding, we estimated a spatial covariance matrix measuring the interdependence between channel pairs, we estimated its position in a Riemannian manifold and we projected it on an Euclidean tangent space obtaining a tangent vector (Fig.1F-G). Finally, we used dimensionality reduction (t-distributed Stochastic Network Embedding, tSNE) to show that tangent vectors that are used as input for covariance-based decoding are linearly separable, while concatenated vectors that are used as input for classic time-domain decoding are not linearly separable (Fig.1E and H). Overall, this simulation revealed that the covariance-based decoding has a specific advantage in decoding temporally misaligned and reciprocally interconnected signals, like the ones driven by cognitive processes such as visual imagery.

### Decoding performance evaluation on MEG data

Then, we used both the covariance-based decoding method and the classic time-domain decoding method to read out imagined categories from MEG signals. Both decoding methods included a single subject level and a group level statistical test. At the single subject level, we computed cross-validated Area Under the Receiver Operating Characteristic Curve (ROC AUC) scores using a sliding window approach. At the group level, we tested whether classification scores were significant across subjects regardless of different amounts of trials used for classification. This second step is necessary because participants presented a different amount of trials after preprocessing (e.g., noise, eye movement, rating trial exclusion). In line with our expectations based on the simulations, the results obtained on empirical data show a stark difference in performance between the two decoding methods. Classic time-domain decoding did not perform significantly above chance level across subjects, in any time window (Fig.2A). In contrast, covariance-based decoding achieved correct classifications significantly above chance level (p<0.05, BF>3) across subjects, in three time windows spanning from 2.5 to 5.5 s (Fig.2B). To test whether these results were determined by the choice of the time window size we also tested shorter time windows. We obtained similar results when using 100 ms time windows for classic time-domain decoding and 500 ms time windows for covariance-based decoding (Fig.2C-D). This suggests that decoding performance is not strictly dependent on the time window size. Even though significant time windows are more sparse when using a shorter sliding window because the temporal misalignment problem is then more pronounced.

**Figure 2.**
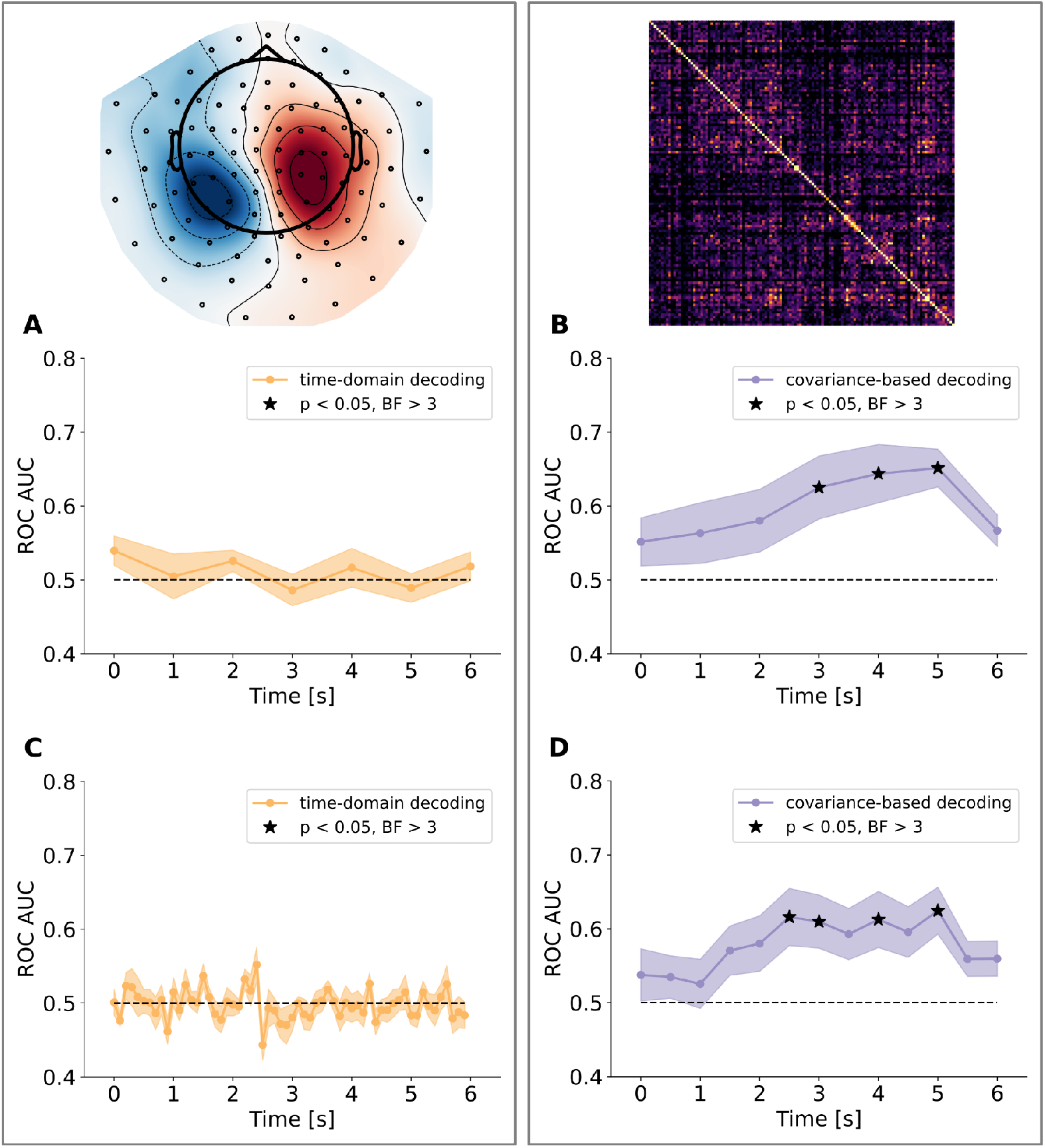
Time-domain decoding and covariance-based decoding applied to visual imagery MEG data. Left panels **(A and C)** show group level decoding results obtained from the classic time-domain decoding method in sensor space (including gradiometers and magnetometers), using 1 s **(A)** or 100 ms **(C)** sliding windows, respectively. Classification score is at chance level and not statistically significant in any section of the trial epoch. Right panels **(B and D)** show Group level decoding results obtained from the covariance-based decoding method, using 1 s **(B)** and 500 ms **(D)** sliding windows, respectively. Classification score is above chance and statistically significant in three (or four) time windows spanning from 2.5 to 5.5 s. The solid line indicates the mean, while the shaded area indicates the standard error of the mean (s.e.m.).

### Relationship between decoding performance and vividness ratings

Next, we tested whether covariance-based decoding performance correlated with participants’ subjective evaluation of the vividness of visual imagery. We tested the relation between decoding performance and subjective ratings both across and within subjects (Fig.3). Across subjects, decoding performance (i.e., ROC AUC scores) correlated with self-reported individual differences in the general vividness of visual imagery (r=0.57, p<0.05, Fig.4A), as assessed by the Vividness of Visual Imagery Questionnaire (VVIQ). Moreover, within subjects, we split the MEG dataset into trials associated with high vividness reports (scores >=3 in the vividness rating provided at the end of the trial, see Fig.1A) and trials associated with low vividness reports (scores<=2), to contrast the decoding performance associated with different subjective vividness ratings. We reasoned that if covariance-based decoding relies on information that contributes to the perceived vividness of visual imagery then high vividness ratings will be associated with higher ROC_AUC scores. Indeed, decoding scores were higher and significant in the time windows from 2.5 to 5.5 s when using only trials with high vividness ratings (>=3, Fig.4B) while the decoding scores were lower and not significantly above chance level (at any time) when using only trials with low vividness ratings (<=2). Importantly, there were no significant differences in vividness ratings between imagination categories (mean face rating = 2.94, mean place rating = 3.01, t = −0.48, p = 0.63) suggesting that participants’ imagination was equally vivid in faces and places trials.

**Figure 3.**
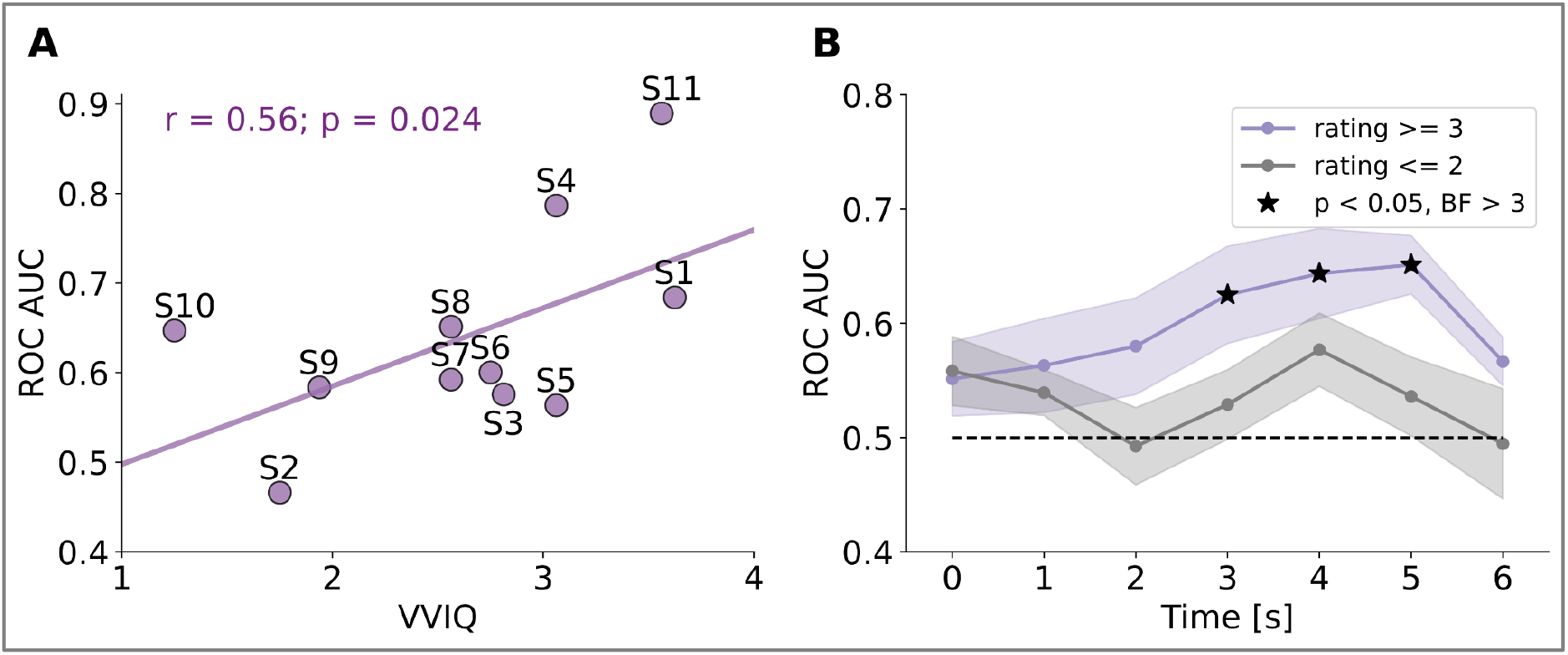
Relation between covariance-based decoding performance and subjective vividness ratings. **(A)** Correlation between vividness of visual imagery questionnaire (VVIQ) scores and ROC_AUC scores obtained using covariance-based decoding. **(B)** We obtained higher and significant covariance-based decoding performance when using only trials associated with high vividness ratings (>=3, purple line), compared to lower and not significant covariance-based decoding performance when using only trials with lower vividness ratings (<=2, gray line). The solid line indicates the mean, while the shaded area indicates the standard error of the mean (s.e.m.).

### Detection and elimination of predictive saccades and microsaccades

In addition, we ran a control analysis to rule out the possibility that the decoding was driven by systematic differences in eye movements associated with face and place imagery. This is a general concern for any neural decoding study, and the covariance-based decoding approach offers a straightforward and clean solution to rule out this potential confound. Although participants were instructed to keep their eyes fixated at the center of the screen during the imagination task, co-registered eye-tracking revealed some residual but systematic micro-saccadic activity that was related to the imagination targets (Fig.4A and B), even after excluding trials with supra-threshold eye-movements (saccades). Therefore, we used covariance-based decoding to read out imagination categories from eye-tracking data that survived the threshold-based exclusion. By visual inspection of eye-tracking data, we observed systematic differences in the eye movement position covariance that may contribute to the classification of face versus place imagery, at least partially for some participants in some time windows (Fig. 4A-B). At the group level, eye movement decoding was statistically significant (p<0.05, BF=3) from 3.5-4.5 s (Fig. 4C). In order to correct for that, we cleaned the MEG dataset from all trials containing any such predictive microsaccades, by training a linear classifier on the eye tracking dataset and estimating the predictive probabilities for each trial. All trials with increased classification probability were excluded from further analyses on the MEG dataset. After predictive microsaccade removal, eye tracking decoding was no longer significant (Fig.4D-F).

**Figure 4.**
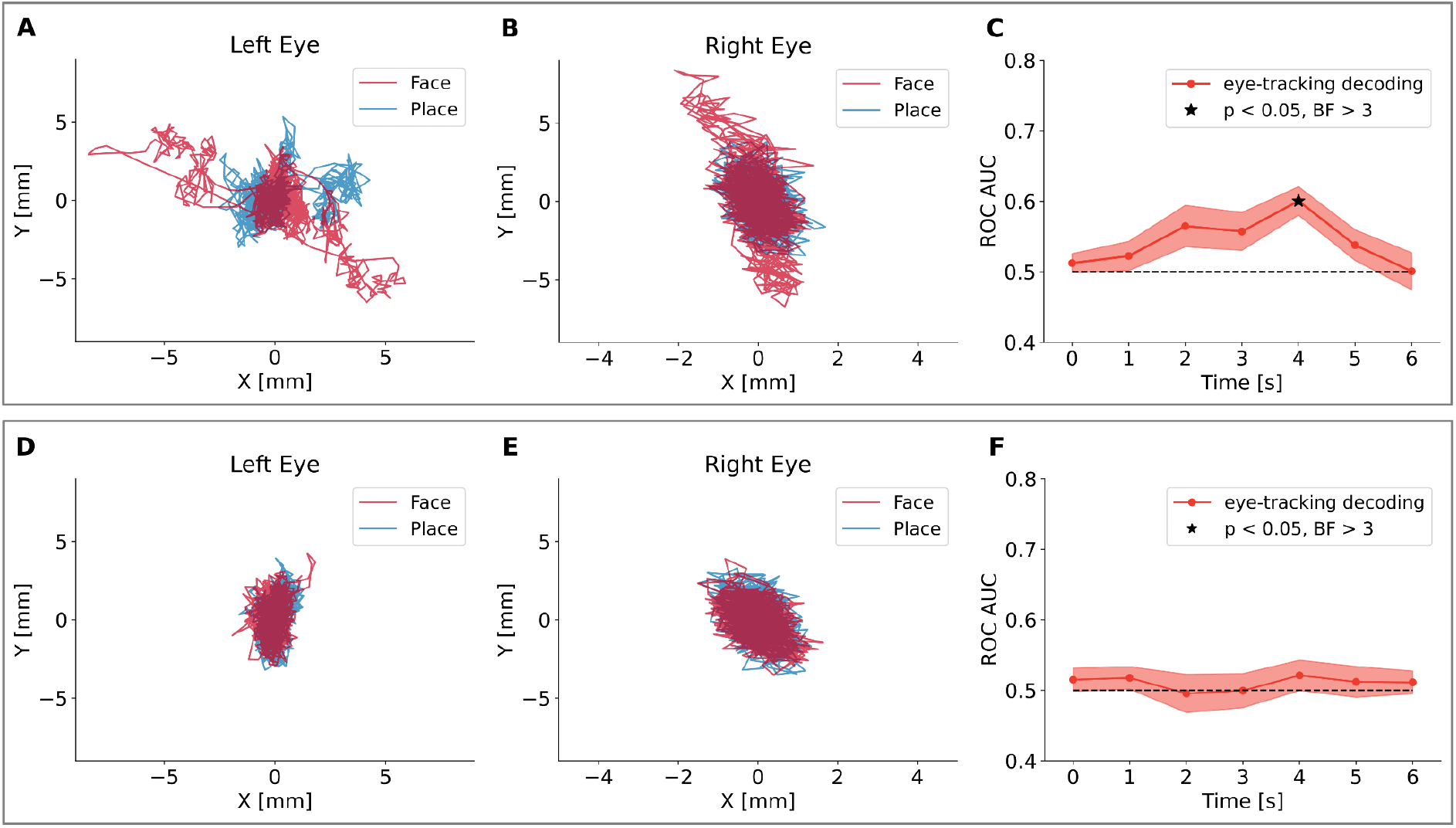
Detection and removal of trials containing predictive microsaccades. **(A, B)** Examples of left and right eye movements in a trial in which the micro-saccade activity contained information about the imagination target (faces, red line, versus places, blue line). **(C)** Group-level decoding based on eye-tracking data only before microsaccade removal (mean and s.e.m). At this level, only trials containing overt saccades exceeding a rejection threshold were excluded. Trials containing sub-threshold, but predictive micro-saccade activation still contributed to the decodability of the imagination target. **(D and F)** After removing all trials with increased decoding probabilities, only trials, in which eye-movement traces did not contain information about the imagination target, remained in the sample. **(F)** Resulting group-level decoding after microsaccade removal.

### The functional connectivity network distinguishing face vs. place imagery

One major advantage of the covariance-based decoding approach for the purpose of this study is that it is inherently based on a functional connectivity measure, i.e. the degree of interaction between node pairs. This allows us to map the most informative connectivity patterns underlying covariance-based decoding. In the following, we use two different types of visualization: edge maps and hub maps (see Fig.5). While the edge map allows us to visualize what sensor pairs are more informative to distinguish face and place imagery, the hub map allows us to visualize what individual nodes are most informative to distinguish face and place imagery. Edge maps are based on the normalized absolute difference in covariance, averaged across trials. Hub maps are based on a cluster-based permutation test between covariance matrices collapsed along one dimension (we refer to this metric as *mstat*, i.e., matrix statistics, for details see Methods).

**Figure 5.**
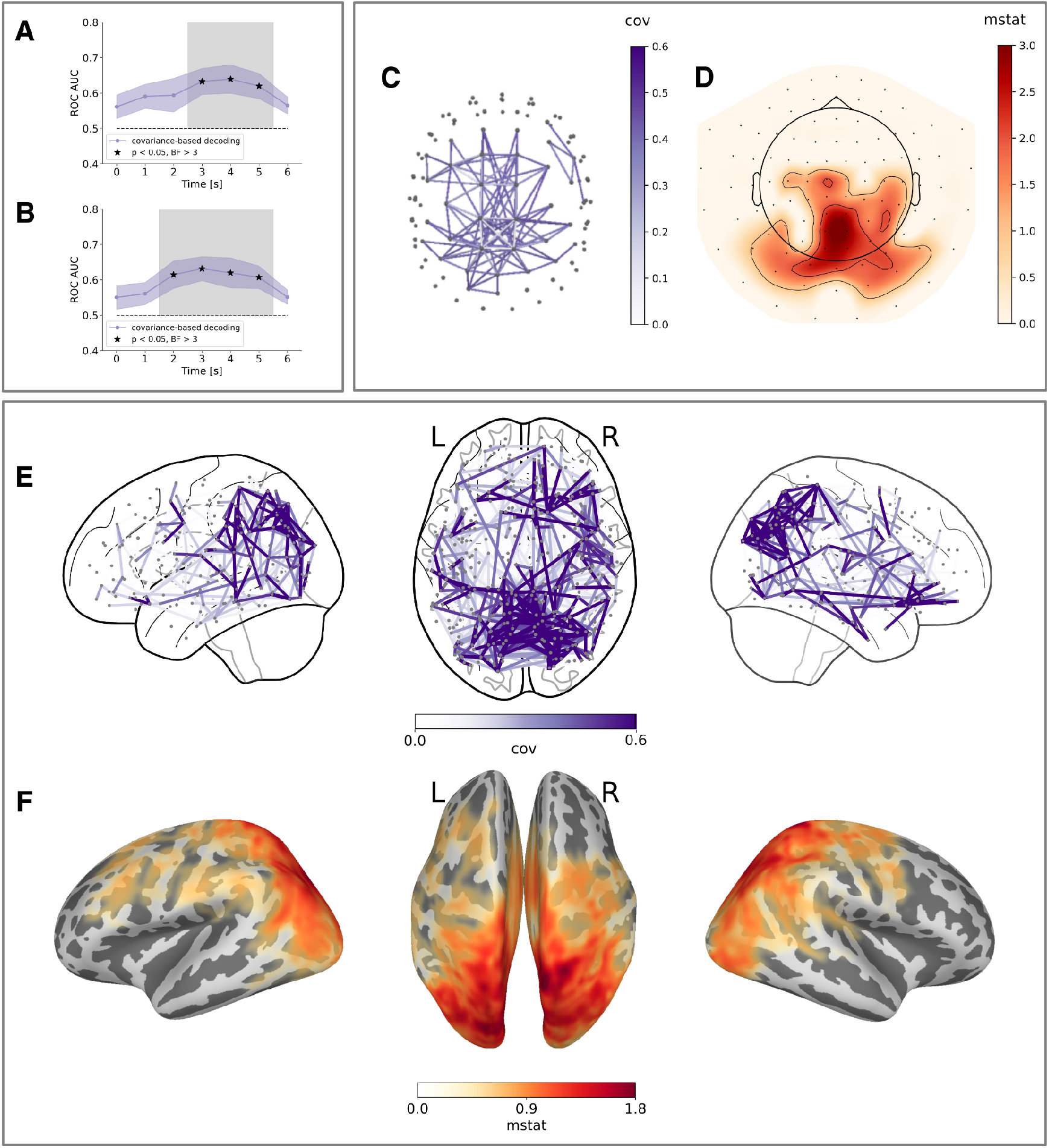
Whole-brain visualization of the connectivity patterns underlying covariance-based decoding in sensor and source space. Decoding results obtained using all sensors (i.e., both gradiometers and magnetometers) **(A)** and all reconstructed sources parcellated using the Glasser atlas **(B),** after removing trials with predictive microsaccades. Edge maps represent the normalized absolute difference in covariance (purple color map). Hub maps represent the output of a cluster-based permutation test between face and place covariance matrices collapsed along one dimension (red color map). **(C)** Edge map showing the most informative connections between sensors distinguishing face and place trials (highest 2 percentiles). Each gray dot represents a sensor and each purple line represents the covariance between two sensors. **(D)** Hub map showing the most informative individual sensors distinguishing face and place trials. **(E)** Edge map showing most informative connections between parcellated areas distinguishing face and place trials (highest 2 percentiles). Each gray dot represents a parcellated area and each purple line represents the covariance between two parcellated areas. **(F)** Hub map showing most informative individual sources distinguishing face and place trials.

To allow for a better localization of these connectivity patterns we applied covariance-based decoding both in sensor space and in source space, which was reconstructed from the MEG recordings using Minimum Norm Estimates (MNE) in combination with 3D models of the subjects’ individual brain anatomies (based on their MRI scans). The reconstructed sources were subsequently parcellated into cortical areas using an atlas. In sensor space, we obtained significant covariance-based decoding (p<0.05, BF>3, Fig. 5A) from 2.5 to 5.5 s using all sensors (i.e., both gradiometers and magnetometers) also after removing trials with predictive microsaccades. The hub map estimated for this time window showed that most informative connectivity hubs distinguishing face and place trials are in the posterior sensors (Fig. 5D). The edge map estimated for this time window - including all connections within the highest 2 percentiles of normalized absolute differences in covariance - showed that the most informative connections include not only short-range connections within both anterior and posterior sensors but also long-range connections between anterior and posterior sensors (Fig.5C). In source space, we obtained significant covariance-based decoding (p<0.05, BF>3, Fig. 5B) from 1.5 to 5.5 s using all parcellated sources also after removing trials with predictive microsaccades. The hub map estimated for this time window showed that the most informative connectivity hubs distinguishing face and place trials are in the occipital and parietal cortices but also in temporal and frontal regions, albeit weaker (Fig. 5F). The edge map estimated for this time window - including all connections within the highest 2 percentiles of normalized absolute differences in covariance - showed that the most informative connections include not only short-range connections within both occipital and parietal areas but also long-range connections between occipital, parietal, temporal and frontal areas (Fig. 5E).

Overall, these results suggest that imagined faces and places involve differences in functional connectivity spanning a broad network of brain areas including not only short-range connections within posterior and anterior areas but also long-range connections between posterior and anterior areas.

### Sub-networks contribution to overall decoding performance

Finally, we tested the contribution from task-relevant sub-networks - including specific regions of interest (ROIs) - to covariance-based decoding. ROIs were selected based on previous literature and included: occipital areas (i.e., dorsal and ventral streams), parietal areas (i.e., inferior and superior parietal), temporal areas (i.e., inferior and medial temporal) and frontal areas (i.e., inferior frontal) (for a complete list see Methods). In particular, the aim of this sub-network analysis was to further disentangle the contribution of short-range and long-range connections to overall decoding performance. By restricting the decoding analysis to subsets of the covariance matrices, we tested the relative contributions of short-range connections (e.g, including distributed nodes within the visual areas) and long-range connections (e.g., including distributed nodes between parietal and visual areas, temporal and visual areas, frontal and visual areas). In this case, since we tested multiple sub-networks at the same time we performed multiple comparisons correction (see Methods). When testing the contribution of short-range connections (Fig. 6A), we obtained significant decoding results (p<0.01, BF>8) using connections within the visual areas (from 2.5 to 5.5 s, light blue line) and within the parietal areas (from 2.5 to 3.5 s, light green line). Decoding within the temporal areas (purple line) and within the frontal areas (orange lines) was not significant. When testing the contribution of long-range connections (Fig. 6B), we obtained significant decoding results (p<0.01, BF>8) between parietal and visual areas (from 1.5 to 3.5 s, cyan line), between temporal and visual areas (from 1.5 to 3.5 s, yellow line), and between frontal and visual areas (from 3.5 to 5.5s, fuchsia line). Decoding between temporal and frontal areas was not significant. We also observed that the decoding was significant when using short-range connections within posterior cingulate areas and long-range connections between posterior cingulate and visual areas (see Fig. S1). This sub-network analysis revealed that both short-range and long-range connections are incremental to overall decoding performance.

**Figure 6.**
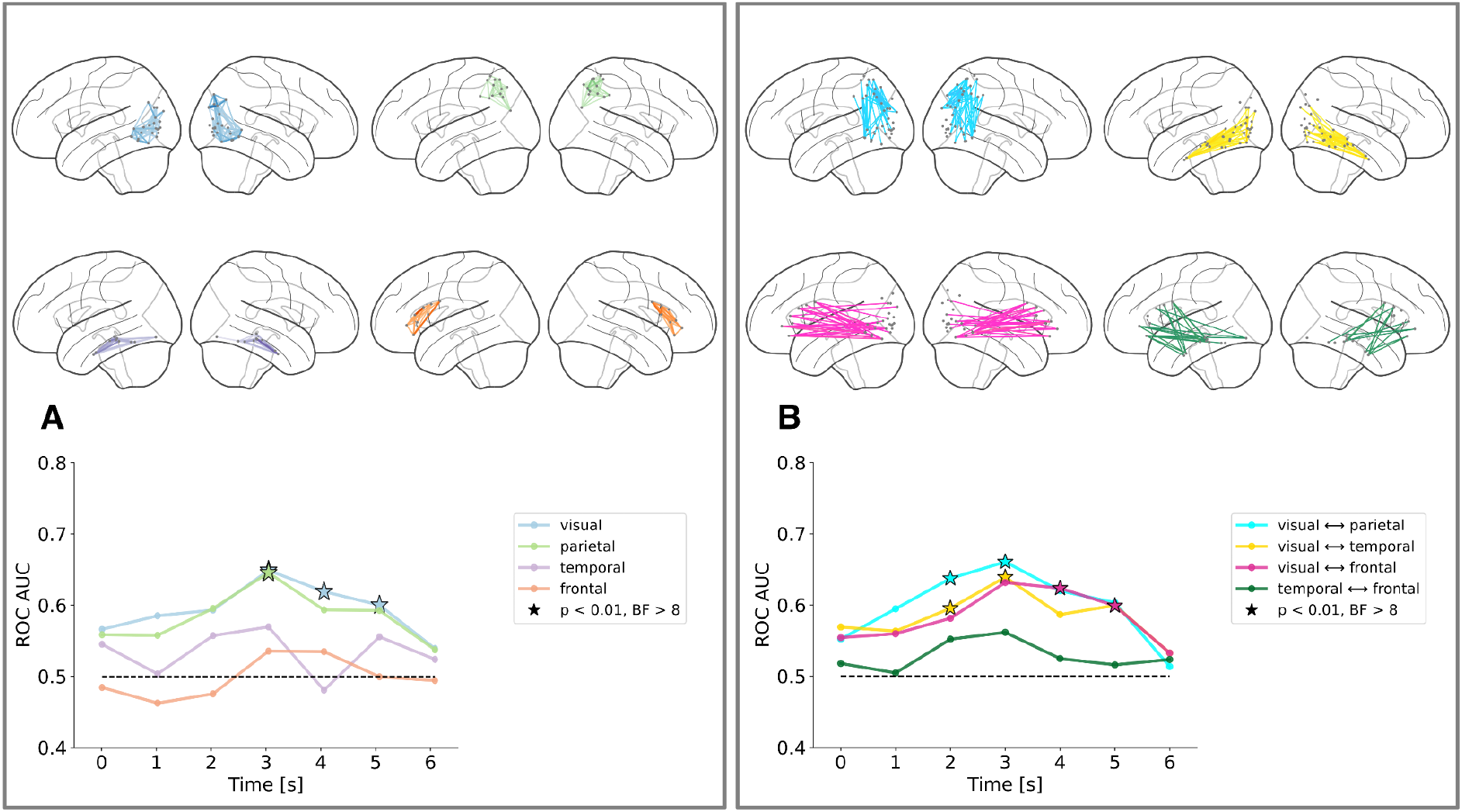
Task-relevant sub-networks contribution to covariance-based decoding. **(A-B)** Task-relevant sub-networks. **(A)** Decoding results obtained using short-range connections within visual (light blue line), parietal (light green line), temporal (purple line) and frontal (orange line) areas. **(B)** Decoding results obtained using long-range connections between visual and parietal areas (cyan line), between visual and temporal areas (yellow line), between visual and frontal areas (fuchsia line) and between temporal and frontal areas (dark green line). For each sub-network, representative connections are shown on a lateral brain view using the same color coding scheme.

To test how spatially specific these contributions from different sub-networks were, we ran a control analysis (Fig. S2) consisting in selecting task-irrelevant sub-networks that were little or not at all involved in the current visual imagery tasks, such as motor areas (i.e., premotor and motor) and auditory areas (i.e., primary and secondary auditory). We observed no significant decoding results for any of the short-range connections within these areas (Fig. S2 A) as well as the long-range connections between these areas (Fig. S2 B). This control analysis revealed that covariance-based decoding relies on spatially specific connectivity patterns associated with task-relevant sub-networks only.

## Discussion

We investigated whether imagined faces and places are associated with distinct connectivity patterns reflecting content-specific feature integration and top-down processing. To do so, we used an experimental paradigm wherein vivid and detailed visual representations were generated internally, in the absence of any external stimulus. Such endogenous neural processes are challenging to decode due to their temporal misalignment across trials and because they involve the cooperation of different brain areas. To address these methodological and theoretical issues we introduced a covariance-based connectivity decoding method, originally designed for brain computer interface (BCI) applications. In particular, we used covariance-based decoding to read out endogenous functional connectivity changes associated with the mental imagery of faces and places. Our results demonstrate the potential and suitability of this decoding approach to answer key questions about the neural mechanisms underlying endogenous brain signals in visual imagery.

One novelty of this study is the use of a minimally constrained experimental paradigm. Previous imagery studies often used a retro-cue paradigm, in which a pictorial cue is displayed on the screen, and participants are instructed to internally recreate that exact mental image, usually in a short amount of time. This paradigm assumes that imagery is similar to visual working memory ^23^, namely the internal replay and maintenance of a recently encountered image. Even though imagery events are better time-locked across trials when triggered by a pictorial cue, there are some methodological issues associated with the retro-cue paradigm. For instance, merely retrieving a recently presented visual stimulus is more constrained and arguably easier as it induces low–level visual features which a decoder can rely on. In contrast, we conceived of mental imagery as a constructive process based on the internal generation of images ^24^ as opposed to a reproductive process based on a replay of recently seen images. Therefore, we did not use any pictorial aid as external reference but word cues only. To preclude the possibility that our decoder could rely on visual signals evoked by visual word cue presentation, the cueing period was separated in time from the subsequent imagination period by a jittered interval of about one second, and furthermore any remaining afterimages were eliminated by a dynamic phase scrambled mask. Moreover, we provided participants with a long imagery time window (6 s), assuming that truly internally re-constructing an image requires time. All these design choices were made to emphasize the internally generated aspects and to minimize potential stimulus-driven aspects. However, the costs of these experimental choices are high in terms of the methodological challenges thereby introduced. In particular, the temporal misalignment of endogenous signals across trials often renders classic time–domain decoding largely ineffective.

We demonstrate a novel methodological approach that can effectively read out purely endogenous signals which, until now, have been challenging to decode from electrophysiological data. Previous studies have investigated the temporal dynamics of visual imagery using time-domain decoding methods that were optimally tuned to decode fast transient changes in brain activity driven by external stimuli ^19^. However, since classic time-domain decoding methods are dramatically impeded by temporal misalignment across trials in mental imagery paradigms, it is necessary to identify methodological solutions. For example, probabilistic decoding models based on latent state dynamics (e.g., Hidden Markov Models) have recently been proposed to deal with the misalignment problem ^25^. The covariance-based decoding approach that we used here is an alternative option to decode temporally misaligned signals. Our simulations showed that temporal misalignment prevents classic time–domain decoding, whereas the covariance–based decoding approach is less susceptible to this challenge and achieves reliable classification across trials. Another important advantage of the covariance-based decoding method is its focus on functional connectivity. Recent fMRI studies using MVPA decoding have shown that imagined categories can be decoded from the sparse co-activation of frontal, parietal and occipital areas ^26^. This was an important step to understand the variety of cognitive processes underlying visual imagery. However, communication across brain areas was not taken into account in this prior work. In contrast, spatial covariance relies on the reciprocal interconnections between brain regions. Decoding methods based on spatial covariance, such as Common Spatial Filter (CSP) ^27^, have been used to decode motor commands from electrophysiological signals for brain computer interface (BCI) applications. This method relies on spatial filters to decode motor commands involving different motor effectors (e.g., left-hand vs. right-hand ^28^). Here, we used an improved version of the CSP method that capitalizes on the geometric properties of spatial covariance matrices.

Importantly, we extend the covariance-based decoding approach to generate both sensor space and source space visualizations of the most informative connectivity patterns, which allows for a meaningful interpretation of contributing hubs and edges all over the cortical fold. In other words, since the signal our decoder is based on directly reflects fluctuations in functional connectivity, we can now pinpoint which functional connections contribute information to solve the cognitive task at hand. The study of functional connectivity networks provides an optimal theoretical framework to answer critical neuroscience questions that are relational in nature ^29, 30^. Measures of statistical interdependence (e.g., correlation) have been previously used to investigate large-scale network dynamics during cognitive tasks. For instance, previous studies successfully decoded different internally driven cognitive states (e.g., free recall, mathematical calculation) from whole-brain connectivity using fMRI ^31^.

In general, the covariance-based decoding method has many advantages including simplicity, interpretability and computational parsimony. The method is mathematically simple because it doesn’t require specific assumptions about frequency and phase, unlike other connectivity measures (e.g., coherence). It is interpretable because it provides information about the connectivity patterns that allows to discriminate between two classes. Information about reciprocal interconnections is also indirectly captured by more complex decoding methods, for instance neural networks. However, neural networks require parameter tuning that is often difficult to interpret. There are also specific types of neural networks models that were specifically designed to capture information encoded in connectivity patterns (i.e., graph neural networks ^32^). However, this network architecture requires a large number of trials which is often prohibitive for neuroscience experiments. In contrast, covariance-based decoding is computationally parsimonious because it requires a smaller number of trials; for instance, here we used 240 trials per participant or less after preprocessing. One limitation is that covariance estimation requires many timepoints to be sufficiently accurate. We obtained significant decoding results using a sliding window of as few as 500 ms. MEG temporal resolution is better than fMRI where covariance estimation would require minutes of task-based recording, which is hardly feasible. However, even covariance-based decoding from MEG might not be enough to decode faster cognitive processes.

To test the validity of the covariance-based decoding method for cognitive neuroscience applications, we performed a series of control analyses to ensure that decoding performance was related to the imagery task and was not driven by potential confounds (e.g. eye movements). A potential confound for every neural decoding study involving a visual task is that classification might be (at least partially) driven by eye movements. There is evidence for the fact that visual imagery, too, may be accompanied by oculomotor activation. For instance, when participants are instructed to imagine a recently seen grid pattern while looking at a blank screen, they produce oculomotor patterns which resemble the oculomotor patterns observed during perception of the actual grid pattern ^33^. Systematic differences in eye movements associated with different imagery categories may affect neural decoding. Previous studies investigating visual perception, visual imagery and visual working memory used co-registered eye-tracking data to rule out potential eye movements confounds ^19, 34^. In contrast, when eye movements are not controlled, there is evidence that they can partially explain neural decoding performance ^35^. The effect of eye movements on neural decoding performance can be explained by the overlap of their underlying neural generators with visual processing. For instance, there is evidence for an extensive overlap between brain areas activated by peripheral oculomotor activity and visual attention ^36^. In order to rule out potential eye movements confounds, we ran a control analysis using co-registered eye-tracking data. We first detected all trials associated with a high predictive probability based on the eye tracking data and then we removed all these trials with predictive eye movements from the MEG dataset. This control analysis is also important for the interpretation of the most informative connectivity patterns associated with covariance-based decoding since it ensures that these connections cannot be explained by systematic differences in eye movements.

Another important step was to show that classification accuracy correlates with participants’ performance in the imagery task. Since there was no direct behavioral measure to assess whether participants performed well or not, we collected subjective vividness ratings for each trial. We expected to obtain better decoding results for those trials in which participants reported having a highly vivid mental image. In line with this conjecture, we observed that higher vividness ratings were associated with higher decoding performance. Moreover, we assessed how well participants considered themselves able to internally generate vivid images in their mind’s eye using the VVIQ. There is evidence for the fact that there are important individual differences in visual imagery ^37, 38^. In particular, the ability to generate mental images ranges from poorly vivid, almost absent imagery (i.e., aphantasia) to highly vivid, almost realistic imagery (hyperphantasia). We expected to obtain better decoding results for participants who considered themselves able to internally generate vivid mental images. In line with this prediction, we observed a significant positive correlation between decoding performance and VVIQ scores (Fig. 3).

Our results have theoretical implications that advance an understanding of the neural mechanisms underlying visual imagery. We asked whether information integration is involved during visual imagery in a way that is similar to visual perception, regardless of whether an external stimulus is presented or not. Since faces and places involve different visual features and different neural generators, we hypothesized that imagining these two different categories will be associated with distinct functional connectivity patterns. In particular, we expected that short-range connections within dorsal and ventral streams - including different brain areas representing specific visual features - will be associated with feature integration during visual imagery. In line with our predictions, we obtained significant decoding results when using short-range connections within visual areas. Our results are consistent with previous studies suggesting that face and scene perception are associated with distinct brain networks ^39, 40^. Indeed, our sub-network analysis included brain areas that are considered to be part of both the face perception network (e.g., the fusiform or occipital face areas) and the scene perception network (e.g., the parahippocampal place area, the retrosplenial cortex, or the occipital place area). We also obtained significant decoding results when using short-range connections within parietal areas. Previous studies have shown that parietal areas are involved in feature integration during visual perception ^41^. There is also evidence from patients with parietal lesions who experience the clinical condition ‘hemispatial neglect’ that is associated with the incapability to visualize a visual hemifield both during perception and imagery ^42^. In line with this literature, we interpret content-specific short-range connections within the parietal areas as reflecting manipulation of spatial information that is necessary to achieve feature integration. These findings, taken together, suggest that information integration does not necessarily require constant monitoring of an external stimulus. In contrast, content-specific coordination between different brain areas associated with feature integration is a basic computation deeply rooted in the nervous system that can be deployed even in the absence of an external stimulus.

A second question we sought to answer was whether top-down processing exerted from cognitive control mechanisms can be specific for different imagination categories. Previous studies have shown local activation during visual imagery in temporal areas associated with memory retrieval ^43, 44^, in parietal areas associated with visuospatial attention ^45, 46^, and in frontal areas associated with focal attention ^11, 47^. These control areas do not only work individually but rather they coordinate with each other in broad networks (e.g., default mode network ^48^ and multiple-demand network ^49^) during cognitive tasks such as memory recall, daydreaming and attentional control. Moreover, there is evidence that, according to the global workspace theory ^50^, control areas constantly exchange information with sensory areas during effortful cognitive tasks ^51^. In line with this view, we expected long-range connections between temporal and visual areas, parietal and visual areas, frontal and visual areas to be associated with top-down processing during imagery. However, it was still unclear whether the contributions from control areas depend on the precise content of imagery or not. We tested whether information contained in long-range connections can be used to distinguish the imagined categories (e.g, face versus place). It is important to point out that we cannot tell apart top-down and bottom-up information flow associated with long-range connections because covariance is a bi-directional functional connectivity measure. Nevertheless, there is evidence suggesting that information is flowing predominantly top-down during imagery and bottom-up during perception ^52, 53^. Our results suggest that long-range connections between temporal and visual areas, parietal and visual areas, as well as frontal and visual areas contain specific information that is captured by covariance-based decoding to distinguish face and place imagery. The specificity of these long-range connections suggests that cognitive control mechanisms do not provide generalized support to visual cortex but rather content-specific information that is tailored to imagined categories (i.e., imagery/attentional templates, see ^14^).

The theoretical and methodological implications of this study extend well-beyond visual imagery. Mental imagery is one example of a purely internally driven cognitive process but there are many others: for instance, endogenous visual attention and visual working memory. All such internally driven cognitive processes are not time-locked to an external stimulus and the endogenous brain activity associated with them is hard to detect because there is no overt behavior. Moreover, internally driven cognitive processes are typically associated with reciprocal interconnections between control areas and perceptual areas. We showed the suitability of the covariance-based decoding approach to answer questions about visual imagery. In addition, we suggest that the same method would be appropriate to also answer research questions about visual attention and visual working memory. Our findings also have implications for visual prediction. Indeed, endogenous signals involving reciprocal interconnections across multiple areas play an important role in visual prediction, as suggested by analysis-by-synthesis ^54^ and predictive coding ^55^ theories. For instance, there is evidence from psychophysics that cueing upcoming images with visual and auditory word cues enhances subsequent visual detection ^56^. However, it is difficult to detect endogenous signals associated with visual prediction. We suggest that the covariance-based decoding approach also offers promising applications in that direction.

Taken together, we showed that imagined faces and places can be decoded from MEG signals using spatial covariance as a measure of functional connectivity. This finding has two implications for the understanding of visual imagery. On the one hand, we show that feature integration in the visual cortex also occurs when there is no external stimulus, and that it is specific to imagery categories. On the other hand, we show that reciprocal interconnections between cognitive control areas and perceptual areas are content-specific, i.e., different for imagined faces and places. To arrive at these conclusions, we used a minimally constrained experimental design that was structured to emphasize the internal generation of mental images. We proposed the application of a covariance-based decoding method originally designed for brain computer interface (BCI) to answer this cognitive neuroscience question. We suggest that our successful application of covariance-based decoding to endogenous signals associated with visual imagery paves the way for future applications to other internally driven cognitive processes, such as visual attention, visual working memory, and visual prediction.

## Materials and Methods

### Participants

Eleven healthy participants (mean age = 28.45, range 24-33, 4 female) with no history of psychiatric or neurological disorders took part in this MEG experiment. All of them reported normal or corrected-to-normal vision. Participants signed an informed consent form before the recording session. Ethical approval to conduct the study was provided by the University of Trento ethical committee.

### Vividness of Visual Imagery Questionnaire

The Vividness of Visual Imagery Questionnaire (VVIQ) is a psychometric test that has been designed to measure individual differences in the vividness of visual imagery ^57^. The VVIQ consists of 16 experimental items organized into four groups. For each group, participants are instructed to imagine a scenario like a familiar person, a familiar shop, or a natural landscape. For each item, participants provide vividness ratings reflecting the visual resolution that they can achieve when they imagine specific details for each scenario (e.g., face contour, characteristic poses, clothes color). Vividness ratings range on a scale from 1 (poor imagination) to 5 (vivid imagination).

Before taking part in the MEG experiment, we asked participants to complete the VVIQ online on an open-source survey platform (LimeSurvey, GmbH, Hamburg, Germany).

### Experimental Procedure

We used the Psychophysics Toolbox ^58^ (PTB-3), MATLAB release R2017b, for stimulus generation and stimulus delivery. The stimuli were projected on a translucent whiteboard using a DLP LED projector (ProPixx, VPixx Technologies Inc., Saint-Bruno, Canada) at a 120 Hz refresh rate. The whiteboard was located at 1 m distance from the participant and it provided a projection area of 51×38 cm (width x height) and 1440×1080 pixel resolution.

The experimental paradigm is shown in Figure 1A. Each trial began with an instruction screen (“Imagine a…”). Then, participants were presented with a visual word cue (“Face” or “Place”) instructing a category for imagination. After that, a fixation cross was shown in the middle of the screen and there was a 600-1600 ms jittered time delay. At this point, the trial epoch started and lasted for 6 seconds. A 15×25 cm picture frame containing a dynamic phase-scrambled mask centered around the fixation cross was displayed on the screen. The picture frame was meant to constrain participants’ imagination to a constant portion of the screen such that the size of the imagined object was consistent across trials and across imagery conditions. Participants were instructed to fill the picture frame with their visual imagination. In particular, they were asked to imagine a familiar face or place of their choosing. Even though participants were allowed to choose the object of their imagination, they were instructed to always imagine the same face and the same place throughout the experiment in order to reduce within-subject variability. Following the trial epoch, participants were asked to rate the vividness of their imagination on a scale from 1 (poor imagination) to 4 (vivid imagination). Finally, we presented participants with a catch question (i.e., “Did you imagine a face or a place?”) in order to make sure they were following the instructions. We used an MEG-compatible response collection system (ResponsePixx Dual Handheld, VPixx Technologies Inc., Saint-Bruno, Canada) to keep track of participants’ responses. Before starting the experiment, participants performed 10 practice trials in order to familiarize themselves with the task. The experiment consisted of 240 trials evenly distributed over 4 blocks. The presentation order of the instructed categories was randomized.

### Data Acquisition

Prior to data acquisition, individual head shapes were digitized with a Polhemus Fastrak digitizer (Polhemus, Vermont, USA), including fiducial landmarks (nasion, right and left pre-auricular points) and about 200 additional points spread out all over the scalp. Five Head Position Indicator (HPI) coils were placed on participant’s mastoid bones and forehead to keep track of participant’s head position inside the dewar through electromagnetic induction before and after each recording block. Landmarks and HPI coils were digitized twice in order to ensure that their spatial accuracy was less than 1mm.

MEG recordings were obtained in a magnetically shielded room (AK3B, Vacuum Schmelze, Hanau, Germany) using a 306-channel (204 first order planar gradiometers, 102 magnetometers) VectorView MEG system (Neuromag, Elekta Inc., Helsinki, Finland). The MEG signal was sampled at 1 kHz, with a low-pass anti-aliasing filter at 330 Hz and a high-pass filter at 0.1 Hz. Before entering the experiment room, we ensured that participants were not wearing or carrying any metallic object and other potential sources of electromagnetic interference. Participants performed the task in a seated position. When positioning participants in the MEG scanner, we ensured tight contact with the dewar. Participants were instructed to avoid head, body and limb movements during the trial epoch.

Moreover, participants were instructed to avoid eye blinks and keep strict eye fixation as much as possible during the trial epoch. Binocular pupil size and eyes’ position were continuously monitored by an MEG-compatible eye-tracking device (Eyelink 1000 Plus, SR-Research Ltd. Mississauga, Ontario, Canada). In the beginning of each experimental session, participants performed an eye-tracking calibration task aimed at verifying the correspondence between pupil position in the image recorded from the camera and gaze position on the screen. Calibration was repeated if drift was noticed in the course of the experimental session.

### Eye-Tracking Analysis

To rule out potential confounds, we removed all trials in which we measured oculomotor noise. In particular, we identified three types of oculomotor noise associated with potential confounds at the brain level: eye blinks, saccades. Eye blinks consist in the rapid opening and closure of the eyelids. Saccades are fast, voluntary eye movements whose amplitude can be up to 15-20 degrees.

We used co-registered eye-tracking data to exclude trials contaminated by oculomotor noise. The eye-tracker measured left and right pupil size (i.e., pupillometry) as well as left and right, horizontal (x) and vertical (y) gaze coordinates. Thus, the eye-tracking output consisted of 6 channels. The analog output was in voltage (−5V to +5V range). The raw eye-tracking signal was sampled at 1 kHz. We segmented eye-tracking data around the trial epoch (0-6 sec). Moreover, we downsampled the raw signal to 250 Hz and applied a notch filter to remove 50 Hz power-line noise. Then, we converted the analog output (in voltage) to digital units (pixels) and we used the physical specificities of the eye tracking device (i.e., data range, voltage range, screen proportion, screen distance) to convert pixels to millimeters. Binocular pupil size was measured in mm^2^. Vertical and horizontal (x, y) binocular gaze coordinates were measured in mm.

For eye blink detection, we used an automatic artifact rejection method based on a pupil size threshold. During blinks the eye-tracking device loses track of the pupil, resulting in missing values in the output file. However, eye blinks are preceded and followed by a sharp decrease in pupil size measurements, because the closure and opening of the eyelids is not instantaneous. For each subject, we computed the absolute value and z-normalized (mean subtracted and divided by standard deviation) pupil area measured from left and right eye. We defined 3 standard deviations from the mean as a threshold for eye blink detection. Trials in which pupil size measurements exceeded the threshold were excluded from further analysis.

For saccade detection, we used an automatic artifact rejection method based on a velocity-threshold identification (VT-I) algorithm ^59^. This algorithm separates fixations and saccades based on their point-to-point velocities using binocular x, y gaze coordinates. We computed the tangent of the rotation angle of the eye relative to the head and we used that measure to calculate eye movement velocities (degrees/second). Velocity profiles typically show two distributions: low velocities for fixations (i.e., <100 deg/sec), and high velocities for saccades (i.e., >300 deg/sec). Trials exceeding the high velocity threshold (i.e., >300 deg/sec) were excluded from further analysis.

Moreover, we tested whether even after we applied the velocity-threshold there were trials containing sub-threshold eye movements (i.e., microsaccades) which were highly predictive for one of the imagination categories. Microsaccades are short-range, involuntary eye movements whose amplitude varies from 2 to 120 arcminutes (1 arcminute = 1/60 of one degree). For predictive microsaccade detection, we ran the covariance-based decoding pipeline (see below) on subthreshold (<300 deg/sec) binocular x, y gaze coordinates. For each trial and each subject, we estimated predictive probabilities for the two imagination categories and we removed all trials with predictive probabilities exceeding a certain threshold. We did not use a fixed probability threshold for every participant, instead we adjusted the probability threshold for each participant (range 65-90%) depending on the average difference in covariance between face and place trials. Trials containing predictive microsaccades in the eye-tracking dataset were excluded from further analysis in the MEG dataset. Finally, we ran the covariance-based decoding pipeline again to test whether predictive microsaccade detection was working properly. To avoid overfitting due to selection bias we used nested cross validation. We divided the eye-tracking dataset in training and test set. We detected trials with predictive microsaccades using the training set and we removed predictive trials from the test set. Then, we divided the portion of data that we previously used as a test set in training and test sets again. We trained the decoding model using the training set and we tested it using the test set. When trials containing microsaccades with high predictive probabilities were removed for each subject, classification scores were at chance level in every time window at the group level (Fig. 4F).

### MEG pre-processing

MEG pre-processing was performed using MNE-Python ^60^ (v0.18.1), Python release 3.6.7, combined with custom routines. First, we removed external and internal sources of noise from the MEG signal. Then, we performed basic signal processing operations like filtering and epoching. External noise (e.g., environmental noise, stationary noise) was removed from MEG recordings offline using a MaxFilter software ^61^ (tsss-filters). In particular, we used a temporally non-extended spatial Signal Source Separation (SSS) algorithm in order to suppress external sources of magnetic interference. Whenever head movements exceed 1 cm within or between blocks, we used the MaxMove algorithm to spatially co-register MEG recordings across blocks to the median head position. HPI movement correction was applied to MEG data collected from 6 over 11 subjects. Then, continuous data was visually inspected for system related artifacts (e.g., SQUID jumps), and contaminated sensors were interpolated. Up to 10 sensors per experimental block were interpolated. Internal noise was reduced using independent component analysis ^62^ (ICA) while preserving signals originating from the brain. Among the potential sources of internal noise there are heartbeat, muscular activity and any residual oculomotor activity (e.g., eye blinks, eye movements) that was not removed based on the eye-tracking data. We used a fixed-point algorithm to estimate 15 independent components in the trial epoch time window (0-6 sec). Up to 5 components per block were excluded based on visual inspection of spatial topographies and latent sources’ time course. A two-pass zero-phase infinite impulse response (IIR) band-pass filter was applied to raw data between 1 and 150 Hz. This IIR filter was based on a Butterworth forward-backward filter. Time series were downsampled to 250 Hz in order to reduce memory load and speed up algebraic operations (e.g., matrix multiplication). Then, we segmented trial epochs from picture frame onset to picture frame offset (0-6 seconds). We further segmented the trial epoch using different time-window segmentation schemas. In particular, we used a short segmentation scheme (100 ms time-windows), an intermediate segmentation scheme (500 ms time-windows) and a long segmentation scheme (1 sec time-windows).

### Trial Exclusion

Trials were excluded from further analyses according to different criteria. We excluded all trials in which the vividness rating was poor (<=2) or participants provided a wrong answer to the catch question about which category they had just imagined. In both cases, the entire trial epoch was discarded. Moreover, we excluded all trials containing oculomotor noise (eye blinks, saccades, predictive microsaccades). In this case, we discarded only noisy time-windows rather than the entire trial epoch. Importantly, the remaining number of trials for each time window was not systematically different between experimental conditions (face vs. place) after trial exclusion.

### Covariance Estimation

We used the pyRiemann toolbox for covariance estimation. We estimated covariance as a measure of joint variability between a pair of time series using Equation 1:

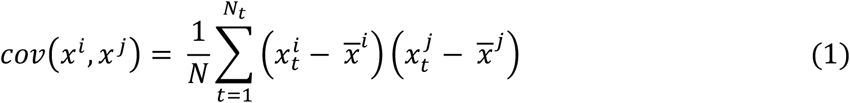

Where x^i^ and x^j^ are time series recorded from different sensors summed across multiple timepoints t divided by the total number of timepoints N.

Spatial covariance matrices (SCMs) were computed as the set of pairwise covariance estimates between all sensors (i.e., 306 × 306 sensors, including both gradiometers and magnetometers), all reconstructed sources (i.e., 5124 × 5124 sources), and all parcellated sources (i.e., 360 × 360 parcels). Covariance estimation can be unstable when the sample size (i.e., trial number) is small and the number of variables (i.e., sensors or sources) is large. Therefore, we used a shrinkage method for covariance estimation ^63^ (OAS) that improves numerical stability and ensures that the matrix is symmetric, positive definite, and thus invertible.

### Data Simulation

Simulated data was generated to compare the performance of different decoding methods using a model of electrophysiological data as close as possible to MEG recordings. We generated time series by summing up three different components. (1) Sinusoidal waves representing endogenous brain signals associated with the experimental task. (2) Band-limited noise representing uncorrelated background brain activity was simulated by summing 50 sinusoids having random frequencies ranging from 1 Hz to 125 Hz, and random phases ranging from 0 to 2π. (3) Pink noise representing the typical 1/f spectral signature of electrophysiological signals was simulated by constructing a power spectral density function for which power is inversely proportional to frequency and applying an inverse fourier transform.

Moreover, we simulated data such that it reflected two main characteristics of endogenous brain signals associated with internally driven cognitive processes, like visual imagery. On the one hand, we added random delays to signal onsets and offsets to account for the fact that endogenous brain signals are not time-locked across trials. On the other hand, We simulated data in 100 trials and 3 recording channels for two different conditions. The two conditions were associated with different spatial configurations that we artificially created by changing the signal to noise (SNR) ratio in the three recording channels. In particular, we simulated data such that the first and the second channel were associated with higher SNR in one condition, while the first and the third channel were associated with higher SNR in another condition.

### Baseline Correction

To rule out the possibility that decoding performance was driven by task-irrelevant individual differences in brain activity and/or brain connectivity we performed a baseline correction. For classic time-domain decoding, the mean of the signal measured in the baseline period was subtracted from the signal measured during the trial epoch. For covariance-based decoding, we used a whitening transformation to remove the covariance measured in the baseline period from the covariance measured during the trial epoch.

### Decoding Analysis

We used two different decoding methods: classic *time-domain decoding* and *covariance-based decoding*. From a methodological point of view, these two decoding methods differ in terms of the brain features used for classification. Time-domain decoding features were obtained by concatenating the raw MEG time-series measured from different sensors into a vector. Covariance-based decoding features were obtained by using a kernel transformation to project spatial covariance matrices from a Rimannian manifold to a locally homeomorphic Euclidean tangent space.

We built a decoding pipeline using scikit-learn toolbox ^64^. This decoding pipeline was applied to MEG data collected from individual subjects. Trial epochs were segmented using a sliding time-window. For each time window, we obtained a classification score. We used three different time-window sizes: 100 ms, 500 ms, 1 s. For time-domain decoding, we standardized the MEG signal by estimating the mean and the standard deviation for each trial and each time-window. For covariance-based decoding, we estimated the spatial covariance matrices for each trial and each time-window. After that, we vectorized our input features following two alternative approaches. For time-domain decoding, we concatenated MEG time series from different recording channels into a single vector for each trial. For covariance-based decoding, we approximated geodesic distances in the Riemannian manifold to Euclidean distances in the tangent space (see Fig. 1G) obtaining a tangent vector for each trial. Then, we used a logistic regression model for binary classification of imagined faces and places trials. In this model, the probabilities of the possible outcomes for each trial are modeled using a logistic function. L2 regularization was applied in order to improve numerical stability. Optimization was performed using a coordinate descent (CD) algorithm that minimizes the cost function by adjusting weights and regularization parameters. Finally, we used the Area Under the Receiver Operating Characteristic Curve (ROC AUC) as a scoring metric. This scoring metric takes into account the tradeoff between true and false positive rates.

### MRI-Based Source Reconstruction

High-resolution T1-weighted anatomical scans were acquired for most of participants (seven over eleven) in a 4T Bruker MedSpec Biospin MR scanner with an 8-channel birdcage head coil (MP-RAGE; 1×1×1 mm; FOV, 256 × 224; 176 slices; TR = 2700 ms; TE = 4.18 ms; inversion time (TI), 1020 ms; 7-degrees flip angle). When the anatomical scans were not available (four over eleven participants) we used a template brain to perform source reconstruction ^65^. This template brain was the average of the anatomical scans collected from 40 subjects (‘fsaverage’). The template brain was deformed to match the headhape of the participants that we measured using the Polhemus Fastrak digitizer (Polhemus, Vermont, USA). For group analysis, we computed a linear interpolation (i.e., morphing) between the individual source model and the template brain for each subject.

The anatomical scans were 3D reconstructed using Freesurfer software ^66^. A Boundary Element Model (BEM) was estimated using the watershed algorithm. MRI and MEG coordinate systems were co-registered by manually matching digitized anatomical fiducial landmarks on the participant’s T1 scan. The resulting whole brain surface reconstruction (5124 vertices; 6.2 mm average source spacing), the BEM model and the aligned coordinate frames were used to compute the 3D forward model for MEG source reconstruction. The inverse operator was estimated using the noise-covariance matrix, the forward solution and the source covariance matrix. We used the Minimum-norm Estimates ^67^ (MNE) for reconstruction of neuronal sources. We used a loose orientation constraint for source reconstruction. In particular, for each source location we estimated a gain matrix having three columns corresponding to magnetic fields *x, y*, and *z* orientations. Then, we computed the norm of these three vectors to obtain one single vector for each source location.

### Cortical Parcellation

To obtain a fine-grained spatial definition of cortical areas and link our results to previous neuroscience literature, we subdivided the reconstructed sources into cortical areas using a multimodal parcellation atlas ^68^. This atlas identifies 360 cortical areas (180 per hemisphere) based on cortical architecture, function, connectivity, and topography. For task-relevant and task-irrelevant sub-network analysis we grouped multiple parcels into larger regions following atlas definitions. Each region included a set of spatially contiguous cortical areas sharing common properties, based on architecture, task-fMRI activity profiles, and functional connectivity. In particular, we selected five larger groups of regions for the task-relevant sub-network analysis: (1) *visual* regions including the following 24 areas for each hemisphere: V1, V2, V3, V3A, V3B, V3CD, V4, V4t, V6A, V7, V8, VMV1, VMV2, VMV3, ProS, PH, FST, IPS1, MST, MT, LO1, LO2, LO3; (2)*parietal* regions including the following 13 areas for each hemisphere: AIP, MIP, VIP, LIPd, LIPv, IP0, IP1, IP2, 7AL, 7Am, 7PC, 7PL, 7Pm; (3) *temporal* regions including the following 10 areas for each hemisphere: EC, FFC, H, PHA1, PHA2, PHA3, PIT, PeEC, PreS, VVC; (4) *frontal* regions including the following 7 areas for each hemisphere: 44, 45, IFJa, IFJp, 47l, IFSp, IFSa, p47r; (5) posterior cingulate regions including the following 7 areas for each hemisphere: DVT, RSC, PCV, POS1, POS2, 7m, v23ab. In addition we selected three larger groups of regions for the tark-irrelevant sub-network analysis: (1) *motor* regions including the following 10 areas for each hemisphere: 4, 55b, 6a, 6d, 6ma, 6mp, 6r, 6v, PEF, SCEF; (2)*primary auditory* regions including the following 5 areas for each hemisphere: A1, LBelt, MBelt, PBelt, RI; (3) *secondary auditory* regions including the following 8 areas for each hemisphere: A4, A5, STGa, STSda, STSdp, STSva, STSvp, TA2.

### Statistical Analysis

Decoding performance was evaluated using statistical tests to establish whether classification was significant both at the single subject level and at the group level.

At the single subject level, we used cross-validation and permutation tests to assess the decoding performance for each time window. In particular, we used a stratified k-fold cross-validation procedure. Data were divided into five folds and classification scores were obtained for each fold. Then, cross-validated decoding performance was estimated by averaging the scores obtained for each fold. Moreover, we ran a permutation test to evaluate the statistical significance of cross-validated scores. This test consisted in repeating the cross-validated classification procedure 1000 times permuting condition labels. We computed the p-value as the percentage of tests for which the classification score obtained with un-permuted labels was greater than the classification score obtained with permuted labels.

At the group level, we evaluated cross-validated decoding performance across multiple subjects using Bayesian hypothesis testing ^69^. To account for the different number of trials per participant resulting from trial exclusion, we used a statistical test that weighs the classification scores for each participant depending on the amount of trials used to train and test the classifier. When we performed more than one test for one single decoding analysis (e.g., sub-network analysis) we corrected for multiple comparisons using False Discovery Rate (FDR) correction.

To investigate which nodes provided most information to covariance-based decoding, we ran a cluster-based permutation test ^70^ (CBPT) both in sensor space and source space. CBPT consists of two different stages: a cluster formation stage and an inferential stage.

In the cluster formation stage, the unit-level statistic is computed for each sensor or source. We used a two-sample covariance matrix ^71^ unit-level statistic that was estimated as follows: first, we estimated the spatial covariance matrices for each trial; then, we computed the element-wise mean covariance matrix and the element-wise variance covariance matrix for each condition; finally, we computed an M (i.e., matrix) standardized statistic that is defined as the squared difference of the mean covariance matrices divided by the sum of the variance covariance matrices. The test statistic is reported in Equation 2:

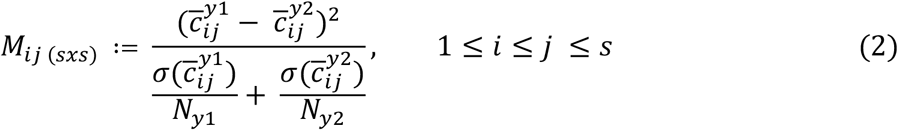

Where M is a sxs (i.e., sensors-by-sensors or sources-by-sources) matrix, c bar is the averaged element-wise covariance estimated between recording channels i and j belonging to either condition y1 or condition y2, and sigma squared is the averaged element-wise variance divided by the number of trials N in each condition. Once we obtained the M matrix, we summed across rows to obtain one single score for each sensor or source measuring the difference in covariance between the two conditions. Given that the distribution of the M standardized statistic is unknown, we run a permutation test under the null hypothesis of exchangeability. We computed the unit-level test statistic 1000 times. For each iteration, assignment to experimental conditions was randomized. Then, the original M values were compared to permuted M values yielding uncorrected p-values. Sensors or sources were selected according to an a priori defined alpha criterion (i.e., p < 0.05) and adjacent sensors or sources not exceeding this value were grouped together into clusters. Finally, we summed all the M values within each cluster (i.e., maxsum) obtaining one single number. Minimum cluster size was set to 5 sensors or 50 vertices. A spatial adjacency matrix containing information about sensors or sources proximity was taken into account in the cluster formation stage.

In the inferential stage, the stored unit-level permutation values summed within clusters were used to compute the cluster-level statistical distribution under the null hypothesis of exchangeability. We calculated the percentage of clusters for which the un-permuted cluster-level statistic was larger than the permuted cluster-level statistic. If the cluster p-value was smaller than 0.05 then we assumed that the data in the two experimental conditions were significantly different.

## Acknowledgments

The authors would like to thank Gianpiero Manitolla and Davide Tabareli for their technical support during data acquisition. The authors also thank Omri Raccah, Arianna Zuanazzi, Joan Orpella and David Poeppel for helpful comments on the manuscript.

## Funding

Fondazione Cassa di Risparmio di Trento e Rovereto (DB)

## Author contributions

Conceptualization: PS, DB
Methodology: FM, EO
Investigation: PS, FM
Visualization: FM
Supervision: EO, DB
Writing—original draft: FM
Writing—review & editing: PS, EO, DB

## Competing interests

All other authors declare they have no competing interests.

## Data and materials availability

All data are available in the main text or the supplementary materials.

## Notes

### Competing Interest Statement

The authors have declared no competing interest.

### Summary of Updates

Title, abstract, results, general format.

