## Supplementary Figures 1-2 for "Covariance-based decoding reveals content-specific feature integration and top-down processing during visual imagery": supplementarymaterials_mantegnaetal2022.pdf

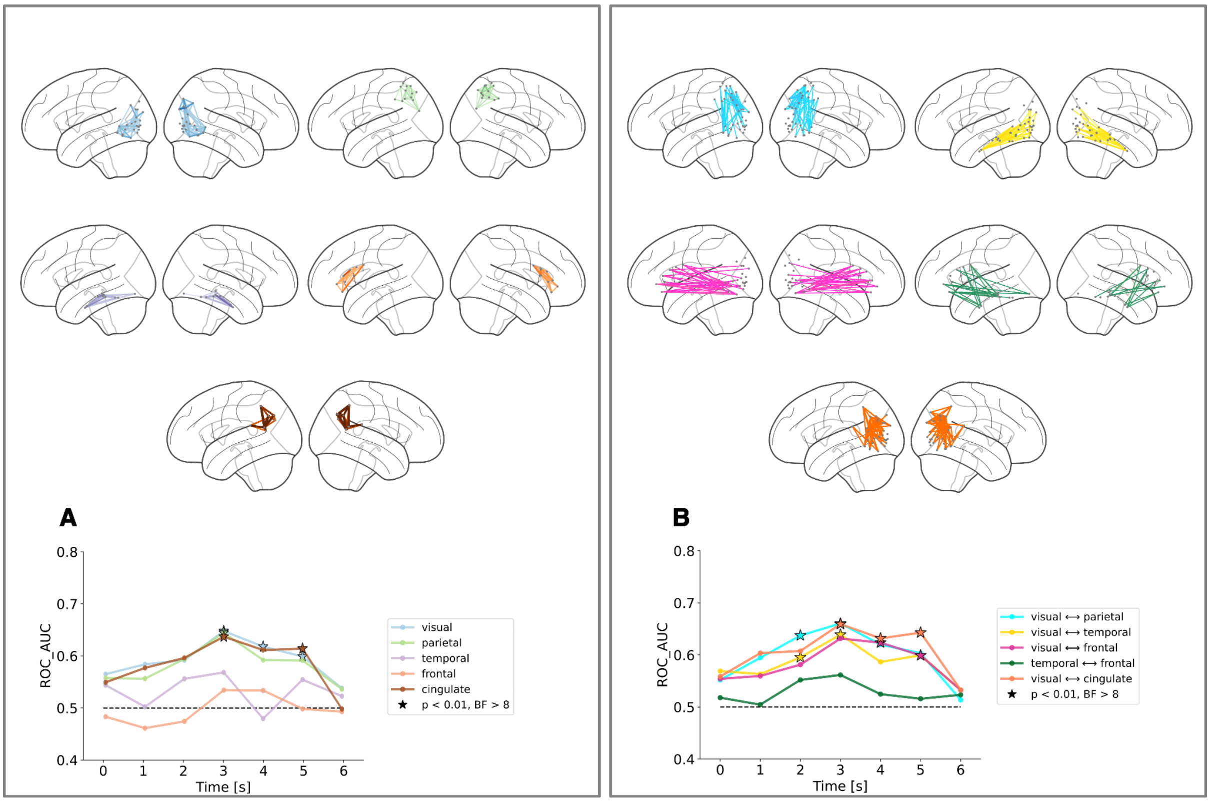


**Figure S1**. **Task-relevant sub-networks contribution to covariance-based decoding**. **(A-B)** Task-relevant sub-networks. **(A)** Decoding results obtained using short-range connections within visual (light blue line), parietal (light green line), posterior cingulate (brown line), temporal (purple line) and frontal (light orange line) areas. **(B)** Decoding results obtained using long-range connections between visual and parietal areas (cyan line), between visual and cingulate areas (orange line), between visual and temporal areas (yellow line), between visual and frontal areas (fuchsia line) and between temporal and frontal areas (dark green line). For each sub-network, representative connections are shown on a lateral brain view using the same color coding scheme.

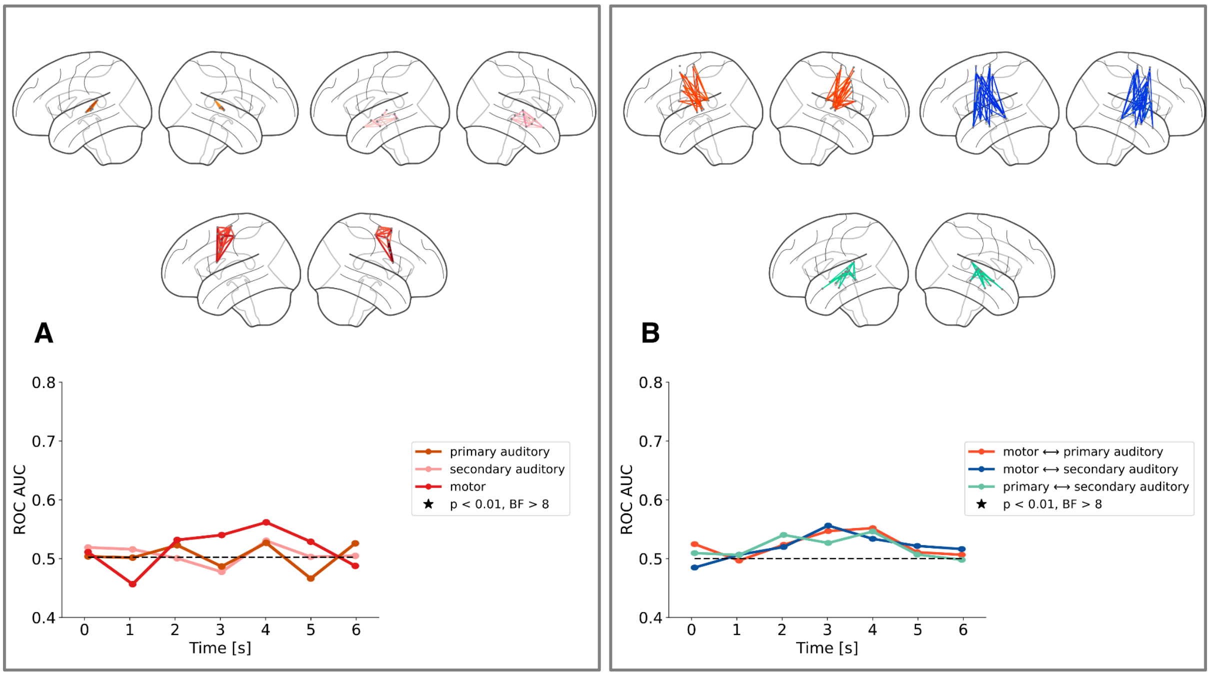

**Figure S2. Task-irrelevant sub-networks contribution to covariance-based decoding**. **(A-B)** Task-irrelevant sub-networks. **(A)** Decoding results obtained using short-range connections within motor (red line), primary (brown line) and secondary (pink line) auditory areas. **(B)** Decoding results obtained using long-range connections between motor and primary auditory areas (red line), between motor and secondary auditory areas (blue line) and between primary and secondary auditory areas (teal line). For each sub-network, representative connections are shown on a lateral brain view using the same color coding scheme.
